## Supplementary information for "Perceptual versus attentional impairments of conscious access: Distinct neural mechanisms despite equal task performance"

This research was supported by a grant from the H2020 European Research Council (ERC STG 715605, SVG).

The experimental blocks of the independent training set differed from the main rapid serial visual presentation (RSVP) task in task context, trial design and stimulus timing (stimuli were presented briefly and in isolation) (**Fig. 1C** vs. **Fig. 3A**) and were collected on a different day, with fewer trials. This led to overall slightly lower decoding performance compared to the main analyses based on training classifiers on the T1 data (**Fig. S2**, top vs. bottom panels), likely reflecting these design differences and greater differences between training and test data with regard to conscious access and working memory demands. Despite these differences, we successfully used these independent classifiers to replicate (**Fig. S6**) and expand upon (**Fig. 3**) the results from our main analyses.

**Figure 3** shows the comparison of T2 illusory triangle decoding between the classifiers trained on the illusory (green lines) and non-illusory triangle in the independent training set (purple lines), because this comparison may isolate illusion-specific processing from basic collinearity-only processing. For this analysis approach to be valid, collinearity-only processing should be similar for the illusory and the non-illusory triangle. To demonstrate that this was indeed the case, we reversed the original cross-feature-decoding scheme: Instead of testing on the illusory triangle, we tested on the non-illusory triangle in T2s, while using the same classifiers that had been trained on either the non-illusory or illusory triangle using the independent training data (**Fig. S7A**, green and purple lines). If collinearity-only processing did not differ between the two types of triangles, there should now be no difference in decoding performance between the classifiers, which is what we found for both the 140-190 ($t_{29}$=-0.90, $P$=0.378, $BF_{01}$=3.56) and the 200-250 ms window ($t_{29}$=0.69, $P$=0.493, $BF_{01}$=4.12; **Fig. S7B**).

We also sought to demonstrate that training on the non-illusory triangle and testing on the illusory triangle in T2s, as reported in the main manuscript, indeed reflected collinearity processing (i.e., the visual properties shared by the two types of triangles, their shape), and no other irrelevant, e.g., task- or attention-related processes. For this purpose, we also trained a classifier on local contrast using the independent training data and tested on the illusory triangle. Local contrast is fully orthogonal to the illusion and has nothing (visual) in common with the illusion, so that any successful decoding would have to reflect non-visual, irrelevant processes. However, we found that this decoding scheme's classifier performance never reached significance ($P$>0.05).

Finally, for local contrast decoding we examined whether the fact that local contrast was not task-relevant or explicitly attended during the main RSVP task could have influenced our findings. We addressed this question by leveraging the manipulation of task-relevance in the independent training set, where either local contrast ("Was the target rotated 180 degrees?"), the non-illusory triangle ("Did the target contain a non-illusory triangle?"), or the illusory triangle ("Did the target contain an illusory triangle?") was task-relevant. For each task-relevance condition, local contrast was decoded using a tenfold cross-validation scheme and the resulting 75-95 ms time window was again averaged (**Fig. S5A**). Surprisingly, we found that classifier performance was somewhat better when local contrast was task-irrelevant ($F_{1,29}$=5.30, $P$=0.008, $BF_{10}$=5.39). However, when we repeated this analysis using data from an independent study with virtually the same task design, we found that task-relevance had no significant effect on local contrast decoding ($F_{1,29}$=1.42, $P$=0.251, $BF_{01}$=3.17; **Fig. S5B**). Local contrast decoding thus appeared largely unaffected by task-relevance.

Using the independent training set, we also replicated the observation that the two consciousness manipulations left local contrast decoding largely intact. For this control analysis, we trained the classifier on the independent training set in which local contrast was task-relevant (**Fig. 3B**, light blue lines). As for our previous main analyses, we averaged the 75-95 ms time window, as this window again contained the decoding peak with occipital topography (**Fig. S2A**, bottom). Similar to our main analyses, another rm ANOVA with the factors masking (present/absent) and T1-T2 lag (short/long) showed that neither masking ($F_{1,29}$=0.97, $P$=0.334, $BF_{01}$=2.90) nor the T1-T2 lag ($F_{1,29}$=0.72, $P$=0.403, $BF_{01}$=3.54) had a significant effect on local contrast decoding, and a paired t-test revealed no evidence of a significant difference between the two performance matched conditions (masked vs. AB condition, $t_{29}$=1.20, $P$=0.240, $BF_{01}$=2.68; **Fig. 3C**, “75-95 ms”).

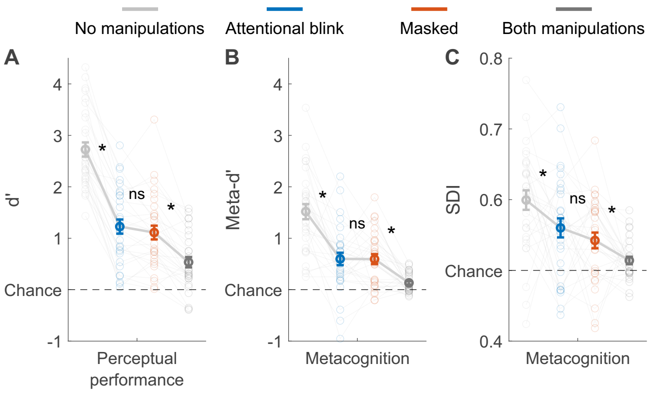


***Figure S1. Alternative measures of behavioral performance.*** *Perceptual performance refers to participants’ ability to detect the Kanizsa illusion. Metacognition refers to participants’ ability to evaluate their own performance using confidence judgments. (A) Perceptual performance in d’. (B) Metacognition in meta-d’* (Maniscalco & Lau, 2012). *(C) Metacognition in subjective discriminability of invisibility (SDI): the area under the receiver operating characteristic curve was calculated after only including trials where participants reported stimulus absence (here, the absence of the illusion). Kanai et al.* (2010) *introduced this measure and found a greater SDI after attentional manipulations compared to perceptual manipulations, suggesting that participants had some knowledge of having missed a stimulus under inattention. However, in the current study, there was no significant difference in SDI between masking and the attentional blink ($t_{29}$=1.17, P=0.250, $BF_{01}$=2.75). Error bars are mean ± standard error of the mean. Individual data points are plotted using low contrast. Ns is not significant (P≥0.176, $BF_{01}$≥2.16). *P≤0.039.*

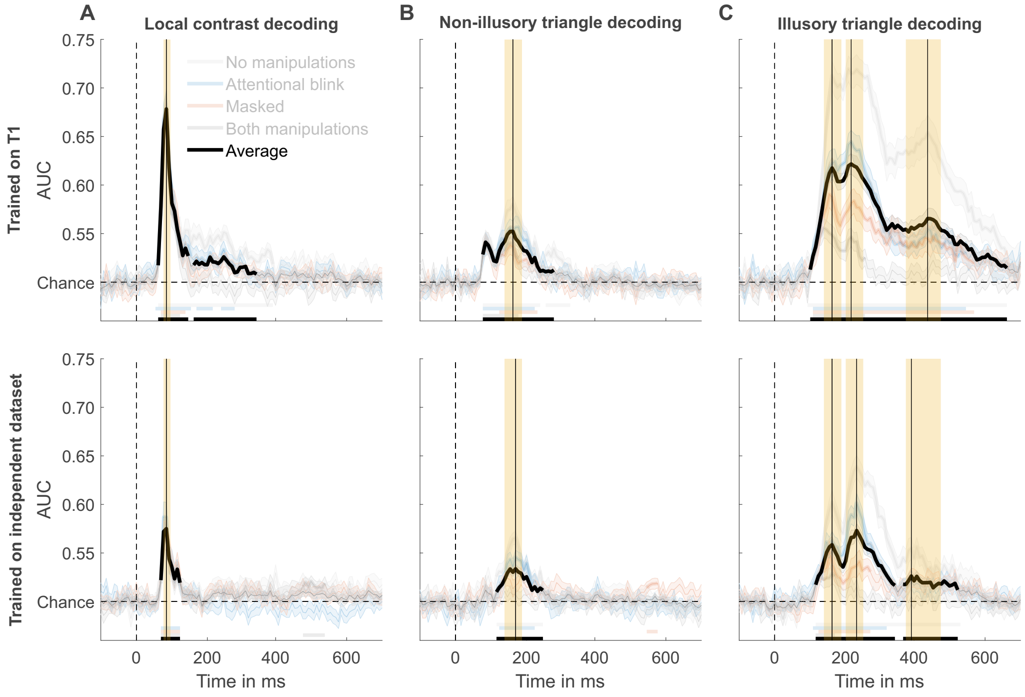


***Figure S2. Averaged decoding accuracy peaks and time window selection.*** *Mean decoding performance, area under the receiver operating characteristic curve (AUC), over time ± standard error of the mean (SEM). Thick lines differ from chance: P<0.05, cluster-based permutation test. Vertical black lines indicate the main peaks and yellow rectangles their encompassing time windows. (A) Local contrast decoding. Regardless of training the classifier on first targets (top) or the independent training data (bottom), the main peak is at 86 ms, which is encompassed by the 75-95 ms time window. (B) Non-illusory triangle decoding. When the classifier was trained on first targets (top), the main peak was at 164 ms. When it was trained on the independent training set (bottom), the main peak was at 172 ms. Both peaks were encompassed by the140-190 ms time window. (C) Illusory triangle decoding. When the classifier was trained on first targets (top), the main peaks were at 164, 219, and 438 ms, which were encompassed by the 140-190, 200-250, 375-475 ms time windows, respectively. When it was trained on the independent training set (bottom), the main peaks were at 164, 234, and 391 ms, which were encompassed by the same time windows.*

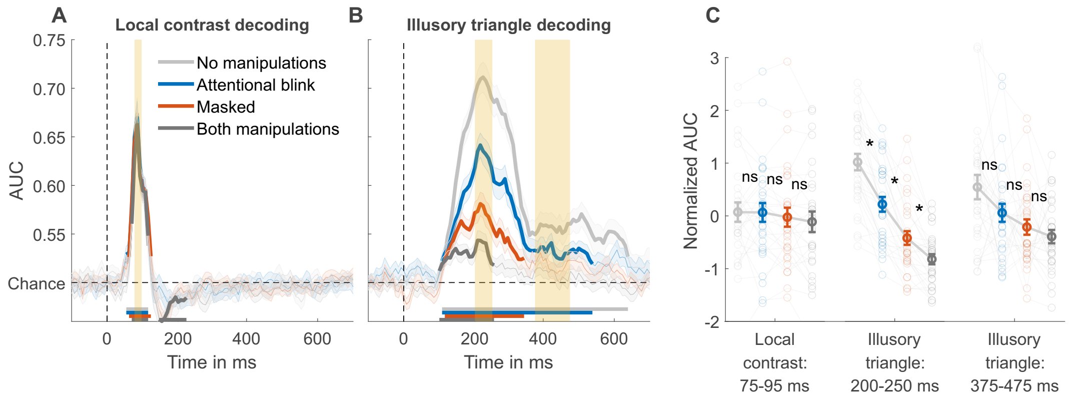


***Figure S3. Local contrast and illusory triangle off-diagonal decoding using first targets as training data.*** *(A) Rotation off-diagonal decoding, trained on the 75-95 ms window of T1. (B) Illusory Kanizsa*

*triangle off-diagonal decoding, trained on the 200-250 ms window of T1. For both features, mean decoding performance, area under the receiver operating characteristic curve (AUC), over time ± standard error of the mean (SEM) is shown. Thick lines differ from chance: $P<0.05$, cluster-based permutation test. (C) Normalized (Z-scored) AUC for every time window. Each window is Z-scored separately. Error bars are mean ± SEM. Individual data points are plotted using low contrast. ns is not significant ($P\geq0.062$, $BF_{01}\geq0.99$). *$P\leq10^{-4}$.*

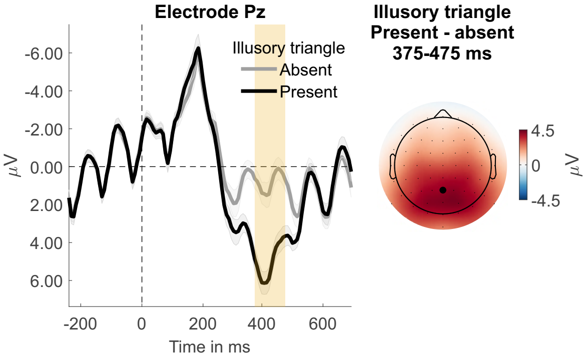


***Figure S4. Event-related potential component P300 derived from the first targets.*** *Electrode Pz waveforms and a topographical voltage distribution are plotted.*

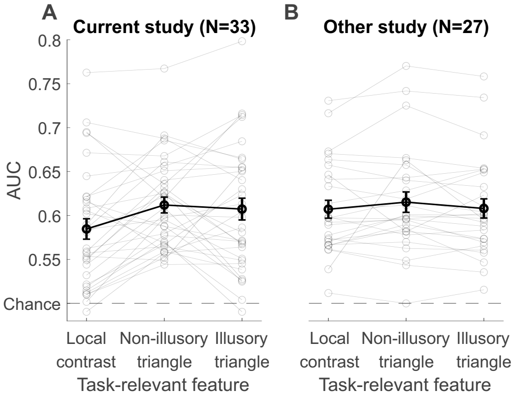


***Figure S5. Local contrast decoding for each task-relevance condition from the independent training data and from another study.*** *Mean decoding performance, area under the receiver operating characteristic curve (AUC), for the 75-95 ms time window. Training and testing were done with the same dataset using a tenfold cross-validation scheme. Error bars are mean ± standard error of the mean. Individual data points are plotted using low contrast. (A) Current study's independent training data (B) Independent study's data.*

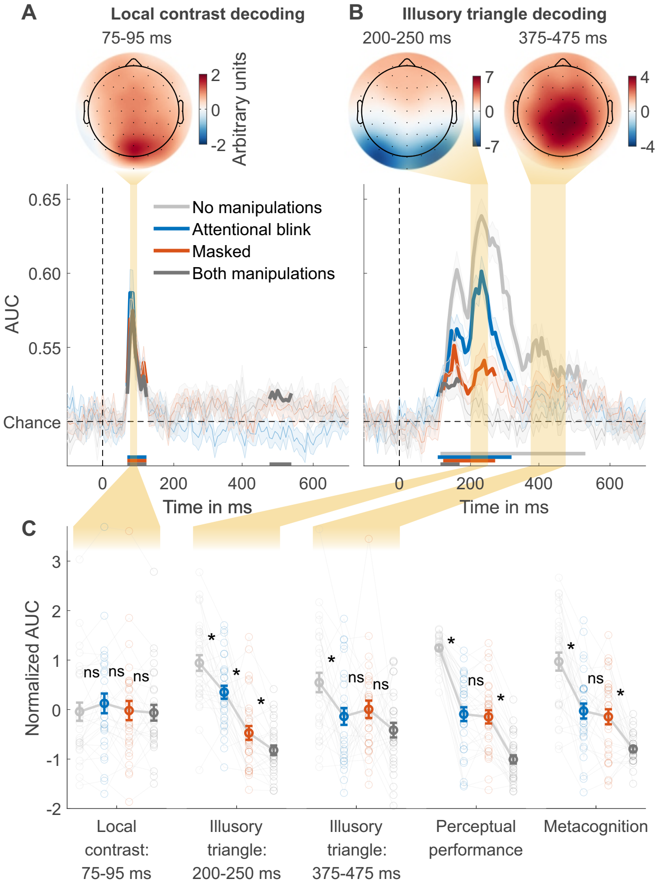


***Figure S6. Local contrast and illusory triangle decoding using the independent training data.*** *(A) Local contrast decoding. (B) Illusory Kanizsa triangle decoding. For both features, covariance/class separability maps reflecting underlying neural sources are shown. Below these maps: mean decoding performance, area under the receiver operating characteristic curve (AUC), over time ± standard error of the mean (SEM). Thick lines differ from chance: P<0.05, cluster-based permutation test. (C) Normalized (Z-scored) AUC for every measure: mean decoding time windows and two types of behavior. Each measure is Z-scored separately. Perceptual performance refers to participants' ability to detect the Kanizsa illusion. Metacognition refers to participants' ability to evaluate their own performance using confidence judgments. Error bars are mean ± SEM. Individual data points are plotted using low contrast. Ns is not significant (P≥0.084,* $BF_{01}$*≥1.26). *P≤0.004.*

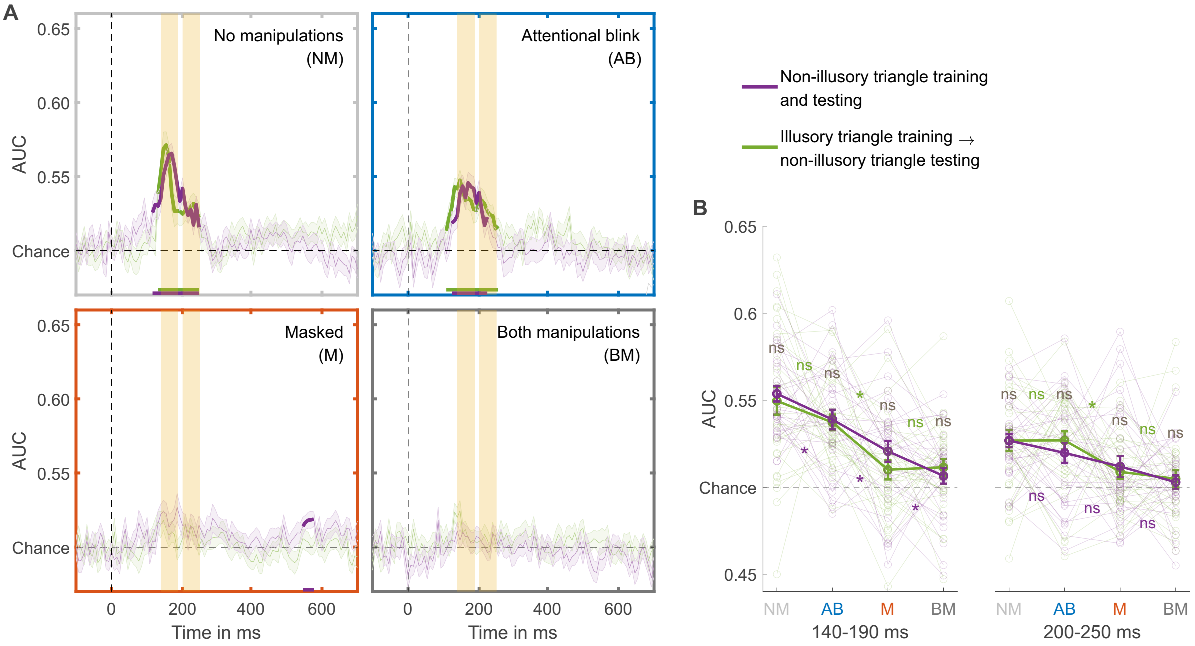


***Figure S7. Control analyses for training on the non-illusory triangle using the independent training set, and testing on the non-illusory triangle from the experimental session.*** *(A) Mean decoding performance, area under the receiver operating characteristic curve (AUC), over time ± standard error of the mean (SEM). The time windows are 75-95 and 140-190 ms. Thick lines differ from chance: $P<0.05$, cluster-based permutation test. (B) Mean AUC for both time windows. Error bars are mean ± SEM. Individual data points are plotted using low contrast. Ns is not significant ($P\geq0.062$, $BF_{01}\geq0.99$). *$P\leq0.040$.*
